# Subacute Inhalation of Ultrafine Particulate Matter Triggers Inflammation Without Altering Amyloid Beta Load in 5xFAD mice

**DOI:** 10.1101/2021.09.07.459017

**Authors:** Liudmila Saveleva, Petra Vartiainen, Veronika Gorova, Sweelin Chew, Irina Belaya, Henna Konttinen, Martina Zucchelli, Paula Korhonen, Emma Kaartinen, Miika Kortelainen, Heikki Lamberg, Olli Sippula, Tarja Malm, Pasi I Jalava, Katja M Kanninen

**Affiliations:** A.I. Virtanen Institute for Molecular Sciences, University of Eastern Finland, Kuopio, 70211, Finland; Department of Environmental and Biological Sciences, University of Eastern Finland, Kuopio, 70211, Finland

**Keywords:** ultrafine particles (UFP), air pollution, 5xFAD, neurodegeneration, neurotoxicology, neuroinflammation

## Abstract

Epidemiological studies reveal that air pollution exposure may exacerbate neurodegeneration. Ultrafine particles (UFPs) are pollutants that remain unregulated in ambient air by environmental agencies. Due to their small size (<100nm), UFPs have the most potential to cross the bodily barriers and thus impact the brain. However, little information exists about how UFPs affect brain function. Alzheimer’s disease (AD) is the most common form of dementia, which has been linked to air pollutant exposure, yet limited information is available on the mechanistic connection between them. This study aims to decipher the effects of UFPs in the brain and periphery using the 5xFAD mouse model of AD. In our study design, AD mice and their wildtype littermates were subjected to 2-weeks inhalation exposure of UFPs in a whole-body chamber. That subacute exposure did not affect the blood-brain barrier integrity or amyloid-beta accumulation. However, when multiple cytokines were analyzed, we found increased levels of proinflammatory cytokines in the brain and periphery, with a predominant alteration of interferon-gamma in response to UFP exposure in both genotypes. Following exposure, mitochondrial superoxide dismutase was significantly upregulated only in the 5xFAD hippocampi, depicting oxidative stress induction in the exposed AD mouse group. These data demonstrate that short-term exposure to inhaled UFPs induces inflammation without affecting amyloid-beta load. This study provides a better understanding of adverse effects caused by short-term UFP exposure in the brain and periphery, also in the context of AD.

**GRAPHICAL ABSTRACT:** 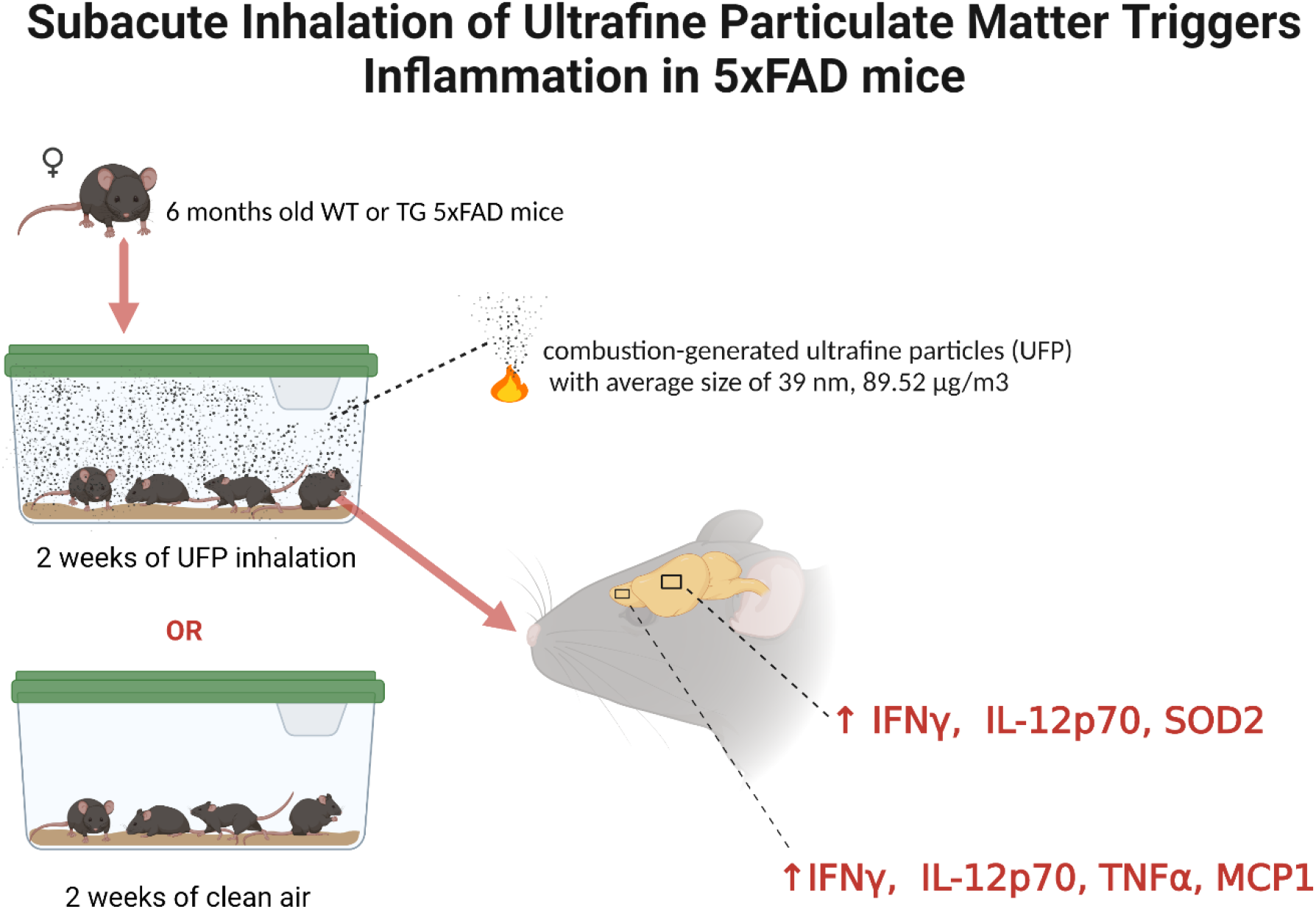

## 1. INTRODUCTION

Air pollution is a serious global concern due to its harmful effect not only on climate but also on human health. The World Health Organization reports that 4.2 million deaths are caused by outdoor air pollution annually. Increasing evidence shows negative effects of air pollutant exposure on respiratory and cardiovascular health, and more recently, reports the association of ambient air pollution to stroke deaths^1,2.^. Recent studies in both humans and preclinical research models demonstrate a link of air pollutants to deteriorating brain health. A systematic review and meta-analysis of epidemiological studies found a positive correlation between air pollution exposure and depression, where residents of highly polluted areas exhibit a higher risk for depression^3^. A correlation between levels of air pollution and the prevalence of mental diseases has also been described^4,5,6^. In rodent studies, long-term exposure to air pollution has been reported to result in impaired or altered behavior, learning and memory disorders, neuroinflammation, oxidative stress, altered morphology of hippocampus, and neuronal loss in the entorhinal cortex^7,8,9,10,11^. Recently, Kim *et al*,. provided a comprehensive review on the impact of fine particulate matter (PM) on neurological disorders, summing up key epidemiological and experimental findings to date^12^.

PM is a key component of air pollution. It consists of solid or liquid aerosol particles suspended in the air, which largely vary in their chemical composition, size, and origin^13^. A commonly used classification of airborne PM is based on their aerodynamic properties. These properties are summarized by the aerodynamic diameter and particles are classified based on diameter range: PM10; <10μm, PM2.5; <2.5 μm, ultrafine PM; <100 nm. Although different components of air pollution may have distinct consequences for health, accumulating evidence suggests that ultrafine PM (UFPs) may have more toxic effects in humans than larger particles^14,13^. It has been shown that UFPs are able to reach the central nervous system via various routes, including crossing both the blood–air barrier, the blood-brain barrier (BBB),^17^ and through the olfactory nerve route^18^. Major sources of anthropogenic outdoor UFPs are industrial processes, road transport emissions, small-scale residential combustion, and power generation^15^. Recently, UFPs have received much attention because of their ubiquitous presence in both indoor and outdoor air, as well as their potential for deeper penetration into the alveolar region and deposition there at high percentages compared to other particles^16^. The toxicity of these particles is heightened due to their small size and large surface area. In our study, we decided to use exclusively the ultrafine fraction of PM due to the previously mentioned toxicity, ability for brain translocation^8^ and neglection of limitation by current policies. We used whole-body inhalation chambers with a relatively low UFP concentration in order not to exceed the annual average PM concentrations reported in the air globally. This allowed us to model more closely the real-life conditions with regular exposure of humans to relatively low doses of UFPs in the ambient air.

Alzheimer’s disease (AD) is the most common neurodegenerative disorder affecting the elderly. Due to the lack of detailed mechanistic insight into the disease cause, current medications only halt disease progression. Environmental factors, such as air pollution, are suggested to contribute to 40% of worldwide dementias and now have been added to the list of the 12 risk factors of dementia^19^. Recent studies have demonstrated that living in highly polluted areas is linked to increased incidence of cognitive impairment and neurodegenerative disease risk, including AD^20,21,22,23.^ Early signs of neurodegeneration are found in the post-mortem brains of adults and children residing in highly polluted areas^24^. Animal studies have shown that prolonged exposure to pollutants can change the inflammatory status of the brain, and promote the development of AD-like pathology^8,25,26,27^. Together, these findings suggest a positive correlation between ambient air quality and neurodegeneration. Particularly, oxidative stress and neuroinflammation are proposed as the main causes of air pollutant-induced adverse effects on the brain^28,29,30,31,14,32,33^. Recent studies have focused on the mechanisms that may contribute to the onset of neurodegenerative diseases following UFP exposure^34,35,36^. However, studies providing insight into how exposure leads to impairments in brain health, and how it exacerbates neurodegeneration, remain limited.

In the current study, we investigated multiple effects of a 2-week UFP exposure in female wild type (WT) and transgenic 5xFAD mice modeling AD, with a special focus on selected brain areas. This mouse strain is widely used not only for studying pathogenesis of AD, but also for testing of therapeutic targets and drugs. Therefore, this AD mouse model was used in hopes of identifying new avenues for investigation of therapeutic targets. In this study design, we used 6-month-old mice to investigate effects of UFP exposure on preexisting AD pathogenesis at an age point when AD mice have already gliosis, amyloid deposition and cognitive impairment. That allowed as to compare the PM exposure effects between AD mice and their WT littermates. We hypothesized that subacute inhalation of UFP will affect oxidative state in the brain and inflammation in both brain and periphery in WT and AD mice, providing putative biomarkers of PM-induced neuroinflammation. This study provides insight into mechanisms altered by UFP exposure in WT mice and mice with pre-existing AD pathology that are affected by a relatively low UFP concentration exposure that closely mimics real-life exposure conditions.

## 2. METHODS

### 2.1. Study design and experimental model

We used female transgenic 5xFAD (*n* =12) mice carrying human amyloid precursor protein (APP) with the APP Swedish, Florida, and London mutations and human presenilin-1 including the M146L and L286V mutations, driven by the mouse Thy1 promoter^37^ and their WT (*n* =10) littermates on the JAXC57BL/6J background. Mice were housed in controlled temperature, humidity, 12:12-h light-dark cycles, and had food and water *ad libitum*. The study was carried out in accordance with the Council of Europe Legislation and Regulation for Animal Protection. The Animal Experiment Board in Finland (Regional State Administrative Agency of Southern Finland) approved all the experiments, which were carried out in accordance with EU Directive 2010/63/EU for animal experiments. The study was conducted and reported according to the ARRIVE guidelines (https://arriveguidelines.org/arrive-guidelines/experimental-procedures).

### 2.2. Inhalation exposures

We used a CAST burner (Combustion Aerosol Standard, Jing Ltd., Switzerland), which is a small, laminar flow, propane-fueled burner, designed for reliable and stable production of soot particles. To prevent complete oxidation of the soot particles, the flame was quenched using nitrogen flow from the side of the burner tip, together with purified dilution air (737-15A, AADCO, USA). The CAST burner has predefined settings to produce soot particles of various sizes. In our experimental setup, the setpoint for particle size was 32nm. The sample was drawn after the burner tip to a porous tube diluter^38^, followed by an ejector diluter (FPS-4000, Dekati Ltd., Finland). The ejector diluter also pushed the sample towards the animal exposure unit. Particle number-size distribution was measured continuously in a scanning mobility particle sizer (TSI Inc., USA), which consisted of a differential mobility analyzer (model 3081, TSI Inc., USA) and a condensation particle counter (model 3776, TSI Inc., USA) utilized in low flow mode (sample flow of 0.3L/min). Furthermore, PM samples were also collected to Teflon filter for gravimetric analysis (MT5, Mettler Toledo), and quartz fiber filter for thermal optical analysis of organic and elemental carbon (OC/EC, Sunset Laboratory Inc.) according to NIOSH 5040 protocol^39^.

When mice were 6 months old, they were randomly divided into four groups: WT-clean air (WT-air, *n* =5), WT-inhalation (WT-UFP, *n* =5), 5xFAD-clean air (AD-air, *n* =6), and 5xFAD-inhalation (AD-UFP, *n* =6). Mice were housed 5-6 per cage, in plastic cages covered with reusable filter animal cage cover (Tecniplast INC., USA) on aspen wood chips, with access to water and maintenance diet *ad libitum* in Scantainer-units. Each exposure group was housed separately. Mouse exposures were conducted in a controlled inhalation whole-body chamber equipped with an automated monitoring system (TSE-Systems, Germany), where CO, CO_2_, and O_2_ concentrations, humidity, temperature, and pressure were monitored^40^. When exposure started, UFP-group mice were exposed in whole-body inhalation chambers in separate inhalation units (5-6 mice per cage, 2 cages per unit) for 5 days/week, 4 h/day, for 2 consecutive weeks. During exposure, mice from the UFP group were housed in two stainless steel cages (TSE-Systems) (L: 265 x W:205 x H:140 mm, 5-6 mice per cage) with access to water and food *ad libitum*. In the exposure chamber, the air exchange was set to 20L/min and the relative humidity to 30%. Clean air for the dilution was obtained from an Aadco 737 zero air generator (Aadco Inst., USA). Mice in the control group received inhalation to the cleanroom air in the same housing conditions. The average UFP particle geometric mean size was 39nm. The average total number concentration was around 7.0×10^5^ particles/cm^3^. The detailed exposure characteristics are presented in Table 1.

**Table 1.** Exposure characteristics.

| Parameter | Value |
| --- | --- |
| The total particle mass concentration ($\mu\text{g}/\text{m}^3$) | 89.5 |
| Number concentration ± SD ($\#/\text{cm}^3$) | $7.0 \times 10^5 \pm 4.9 \times 10^5$ |
| Geometric mean size ± SD (nm) | $38.54 \pm 8.22$ |
| Organic carbon content ($\mu\text{g}/\text{m}^3$) | 17.3 |
| Elemental carbon content ($\mu\text{g}/\text{m}^3$) | 25.9 |
| Total carbon ($\mu\text{g}/\text{m}^3$) | 43.2 |

All mice were weighed 1 day before inhalation started, 1 week after inhalation exposure, and on the day of sacrifice. There were no statistically significant differences in body weights between mouse genotypes, or after the 2 weeks of UFP exposure (data not shown).

### 2.3. Tissue collection

After 2-week exposure, the mice were terminally anesthetized with 2,2,2-Tribromoethanol (Sigma-Aldrich, USA), followed by transcardial perfusion with heparinized (2500 IU/L, LEO) 0.9% saline before organ collection. Blood was collected through cardiac puncture and centrifuged at 1300*g* for 10min at 4°C to isolate plasma. Plasma samples were stored at −70°C until further use.

The brains were dissected out of the skull, cut midsagittally. The left hemisphere was further processed for immunohistochemistry, while the right hemisphere was collected for biochemical analyses. The left hemisphere was immersion fixed with 4% paraformaldehyde in 0.1M phosphate buffer (pH 7.4) for 22h. Then the left hemisphere was incubated in 30% sucrose solution for 48h, and later frozen in liquid nitrogen and stored at − 70°C until cryosectioning. For the right hemisphere, olfactory bulbs and hippocampi were isolated and snap-frozen in liquid nitrogen and stored at −70°C until protein and RNA isolation.

### 2.4. Protein and RNA isolation from brain samples

Snap-frozen brain samples were homogenized manually in eight volumes of lysis buffer, pH 7.4, containing 20mM Tris, 250mM sucrose, 0.5mM EDTA, 0.5mM EGTA, 4% v/v protease, and 1% v/v phosphatase inhibitor cocktails (Sigma-Aldrich, USA). Cytosolic extracts from the tissues were collected by centrifugation at 5000*g* for 10 min at + 4 °C. The supernatants were aliquoted and stored at − 70°C. Pierce 660nm Protein Assay Kit (Thermo Fisher Scientific, USA) was used to determine the protein concentrations. One of the sample aliquots was mixed with TRI Reagent (Sigma-Aldrich, USA) for total RNA isolation following the manufacturer’s instructions.

NanoDrop 2000 (Thermo Fisher Scientific, USA) was used for determining the concentration and purity of RNA. RNA samples with 260/280 ratios higher than 1.8 were used for analyses.

### 2.5. Quantitative real-time PCR

Genomic DNA was removed from all RNA samples with DNase I, RNase-free kit (Thermo Fisher Scientific, USA) according to the manufacturer’s protocol. 1μg of mRNA was reverse-transcribed using the High-Capacity cDNA Reverse Transcription Kit (Thermo Fisher Scientific, USA). Quantitative real-time PCR (RT-PCR) was performed by using StepOnePlus Real-Time PCR System (Thermo Fisher Scientific, USA). All TaqMan gene expression assays (Thermo Fisher Scientific, USA) used for RT-PCR in the study are listed in Supplementary Table 1. Ct values were normalized to the glyceraldehyde 3-phosphate dehydrogenase (*Gapdh*) endogenous control. Fold changes of gene expression were calculated using the ddCt method.

### 2.6. Western blot

20μg of protein lysate from hippocampal samples was used for Western blot. Proteins were separated on 12% SDS-PAGE gels and transferred to polyvinylidene difluoride membranes (Cytiva Amersham™ Hybond™, USA) with Trans-Blot Turbo Transfer System (BioRad, USA). The membranes were blocked in 5% fat-free milk in phosphate-buffered saline containing 0.2% of Tween-20 (PBST) before incubation with primary antibodies overnight at 4°C. After incubation, the membranes were then washed thrice with PBST and incubated with secondary antibodies for 2h at room temperature. For HRP-conjugated antibodies, the SuperSignal™ West Pico PLUS Chemiluminescent substrate kit (Thermo Fisher Scientific, USA) was used for the visualization of bands. The BioRad ChemiDocTM Imaging System was used for imaging. The amount of protein was quantified by using ImageLab software (BioRad, USA). All antibodies used are listed in Supplementary Table 2.

### 2.7. Cytokine bead array

The cytokine bead array mouse inflammation kit (BD Biosciences, USA) was used for the determination of cytokine levels in brain lysate and plasma samples according to the manufacturer’s protocol. 15μg of hippocampal and olfactory bulb protein lysates or proteins from plasma were used to measure concentrations of the following cytokines: including tumor necrosis factor alpha (TNFα), interferon gamma (IFNγ), monocyte chemoattractant protein 1 (MCP-1), interleukin 6 (IL-6), IL-10, IL12p70. CytoFlex S flow cytometer (Beckman Coulter, USA) was used for the measurement, and acquired results were exported and analyzed with FCAP Array v3.0 software (Soft Flow Inc., USA).

### 2.8. Immunohistochemistry

Fixed brains were cut in serial 20μm sagittal sections, each section 240μm apart, using a cryostat (Leica Microsystems, Germany), and stored submerged in anti-freeze solution at − 20°C. Before staining, sections were washed in 0.1M PBS and placed on super frost glass slides (6 sections/slide). To prevent unspecific binding, sections were blocked for 1h in 10% normal goat serum in PBST and then incubated overnight at room temperature with primary antibodies. The following day, sections were washed in PBST and incubated for 2h with suitable fluorescent secondary antibodies diluted in 5% normal goat serum in PBST. All primary and secondary antibodies are listed in Supplementary Table 2. A total of 3-6 sections 240μm apart were imaged for each animal. Images were taken with 10x magnification on the Axio Imager M2 using a digital camera (Axiocam, Zeiss, Germany) and ZEN software. Each image was stitched to avoid overlap of adjacent tiles and exported in TIFF format. The signal was quantified using Fiji software. The percentage of immunopositive cells or positively stained area (in case of extracellular localization of antigen) within the region of interest was measured for each brain section, and the average from 3-6 sections per mouse was reported.

### 2.9. Statistical analyses

Statistical analyses were performed with GraphPad Prism version 8.0.2 for Windows (GraphPad Software, USA) and ARTool-package for R^41^. The statistical outliers were identified with Grubbs’ test and removed from the analysis. Data were tested against the null hypothesis that it is normally distributed. Two-way analysis of variance (ANOVA) was used to examine the main genotype, UFP-inhalation exposure, and genotype*UFP interaction effect between WT and 5xFAD mice in case of normal distribution. This followed by an unpaired *t* test comparison in case of a significant interaction between two factors (genotype*inhalation) to examine inhalation effect for each group separately. For a nonparametric data, nonparametric ANOVA test was used. In case of a significant interaction between two factors (genotype*inhalation), the Mann-Whitney test was used to examine the inhalation effect within one genotype. All values are expressed as mean ± standard error of the mean (SEM). A difference was considered significant when *p* <0.05.

## 3. RESULTS

### 3.1. Subacute UFP inhalation did not affect amyloid beta burden or blood brain barrier integrity in exposed mice

The brain amyloid-beta (Aβ) burden has been shown to be increased upon air pollutant exposure in humans^20^ and rodents^8,34,42^. Therefore, we first assessed whether the levels of Aβ were altered in the brains of WT and 5xFAD mice following a 2-week UFP inhalation exposure. Histochemical staining with the Aβ antibody WO2 revealed the presence of plaques in hippocampal and cortical brain areas of 5xFAD mice, as expected (Supplementary Fig. 1). However, a 2-week UFP inhalation exposure did not affect the levels of Aβ in the hippocampi or cortex of the 5xFAD mice (Supplementary Fig. 1). To study whether UFP exposure impacts the permeability of the BBB, we analyzed the levels of Glucose transporter 1 (GLUT-1) by immunohistochemical staining, as previously reported^43^. UFP inhalation exposure did not alter GLUT-1 levels (Supplementary Fig. 2A), suggesting that the BBB was not compromised due to the sub-acute inhalation exposure. Furthermore, exposure did not alter the mRNA expression level of Gap junction alpha-1 protein (GJA1) in the olfactory bulbs and hippocampi of both WT and 5xFAD mice (Supplementary Fig. 2B, C), further corroborating our finding that the BBB remains intact upon exposure.

### 3.2. Systemic inflammatory responses to UFP exposure

Given that UFPs have been reported to influence inflammatory processes^28,31,29,32,44^ and that increased inflammatory cytokines in plasma were associated with severity of AD in humans^45,46^, we next deciphered the impact of UFP exposure on systemic inflammation using plasma samples from exposed mice. CBA analysis revealed that interferon-gamma (IFNγ) was significantly up-regulated in the plasma after UFP-inhalation in both WT (18.16 ± 0.49pg/ml in WT-UFP group compared to 15.96 ± 0.88pg/ml for the WT-air group) and 5xFAD mice (17.71 ± 0.34pg/ml in AD-UFP group compared to 17.08 ± 0.52pg/ml for AD-air group) (p < 0.01) (Fig. 1A). interleukin 12p70 (IL-12p70) was significantly increased in plasma of AD-UFP group (35.97 ± 4.33pg/ml) compared to AD-air group (15.33 ± 4.68pg/ml, p < 0.05) (Fig. 1B). Interestingly, Pearson correlation analysis revealed a significant (p < 0.05) positive correlation between plasma and hippocampal concentrations of IFNγ in the AD group (r = 0.67), indicating similar responses to UFP exposure in this group in both brain and periphery. Other measured cytokines remained unaltered in the plasma of both WT and 5xFAD mice following exposure (Supplementary Fig. 3).

**Fig. 1.**
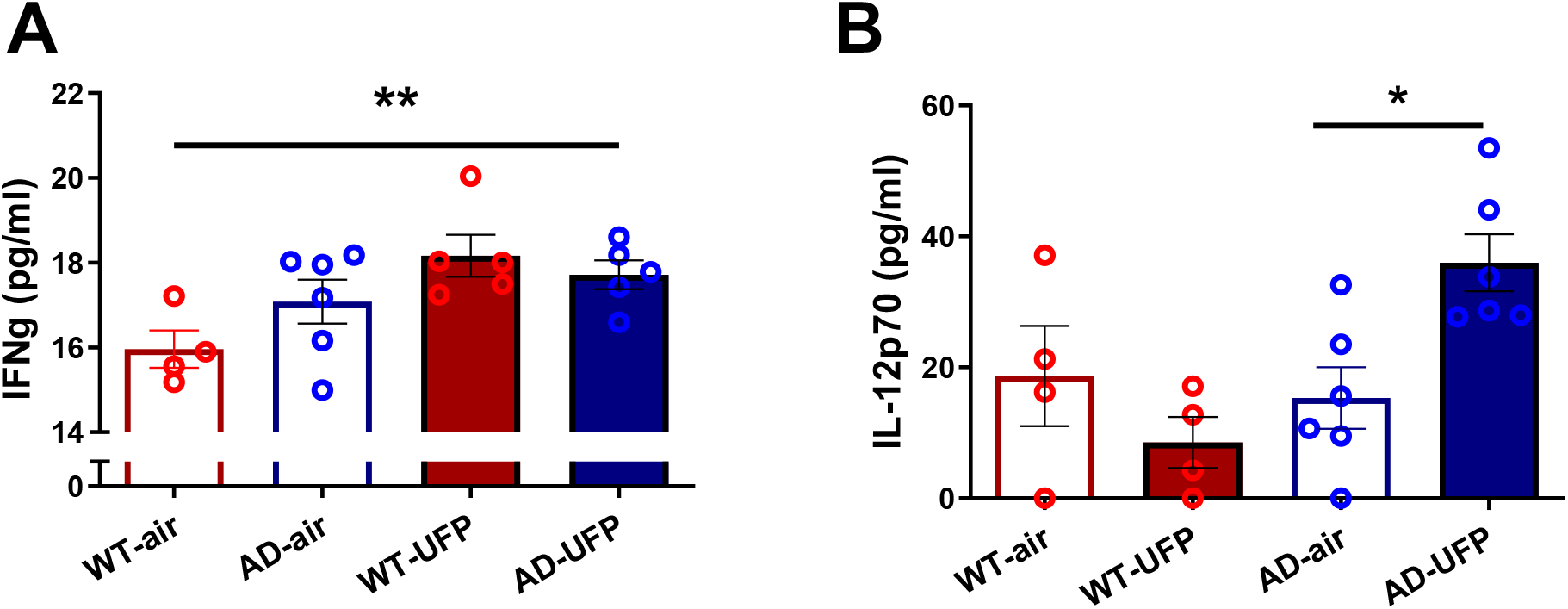
A Increased IFNγ in plasma after 2-week UFP inhalation exposure. All data are presented as mean (±SEM). Two-way ANOVA was used to measure genotype and UFP exposure effect between WT and 5xFAD mice. **B**. Upregulated IL-12p70 in plasma of AD mice after UFP exposure. Nonparametric two-way ANOVA was used to measure the genotype and UFP exposure effect between WT and 5xFAD mice. Because of a significant genotype*UFP interaction, the Mann-Whitney test was used for the comparison of groups within one genotype. «*» UFP inhalation effect: *p < 0.05, **p < 0.01.

The basal cytokine levels of non-exposed mice (WT-air vs. AD-air) were not altered due to mouse genotype at the age of 6 months.

### 3.3. Inflammatory responses to UFPs in olfactory bulbs

To investigate the effect of UFPs on one of the proposed entry gateways of particles to the brain, we next measured cytokine levels in the olfactory bulbs. UFP-induced neuroinflammation was observed regardless of genotype, in both WT and 5xFAD mice, as evidenced by alterations to tumor necrosis factor-alpha (TNFα), monocyte chemoattractant protein 1 (MCP-1), IFNγ, and IL-12p70 (Fig. 2A-D). Significantly higher levels of TNFα were observed in WT-UFP mice (21.10 ± 0.82pg/ml) when compared to WT-air mice (18.12 ± 0.24pg/ml, p < 0.001). Likewise, TNFα level in AD-UFP (20.55 ± 0.74pg/ml) was higher than AD-air group (18.10 ± 0.32pg/ml, p < 0.001). MCP-1 level was higher in WT-UFP group (202.0 ± 10.1pg/ml) than WT-air group (119.8 ± 9.80pg/ml) and higher in AD-UFP group (116.1 ± 14.97pg/ml) than AD-air group (70.93 ± 19.57pg/ml), (p <0.0001). In addition, we observed a genotype-dependent difference in MCP-1 levels (Fig. 2B). IFNγ level was higher in WT-UFP group (17.46 ± 0.64pg/ml) than WT-air group (16.35 ± 0.13pg/ml, p < 0.01). Similarly, IFNγ level was higher in AD-UFP group (16.99 ± 0.46pg/ml) than AD-air group (15.44 ± 0.32pg/ml, p < 0.01). IL-12p70 levels increased in WT-UFP group (25.53 ± 5.53pg/ml), when compared to WT-air group (8.95 ± 3.10pg/ml), and also in AD-UFP group (21.67 ± 4.85pg/ml), when compared to AD-air group (16.56 ± 3.38pg/ml) (p < 0.05). IL-10 level in AD-UFP (39.88 ± 1.76pg/ml) was lower than WT-UFP mice (56.41 ± 5.35pg/ml, p < 0.05), supporting a genotype-dependent effect. Interestingly, for most of the cytokines (TNFα, IL-12p70, IL-10) we observed a higher magnitude of response to UFP inhalation in WT mice than in AD mice. UFP exposure or genotype-specific changes were not detected in IL-6 in the olfactory bulbs of both WT and 5xFAD mice in our experiment (data not shown).

**Fig. 2.**
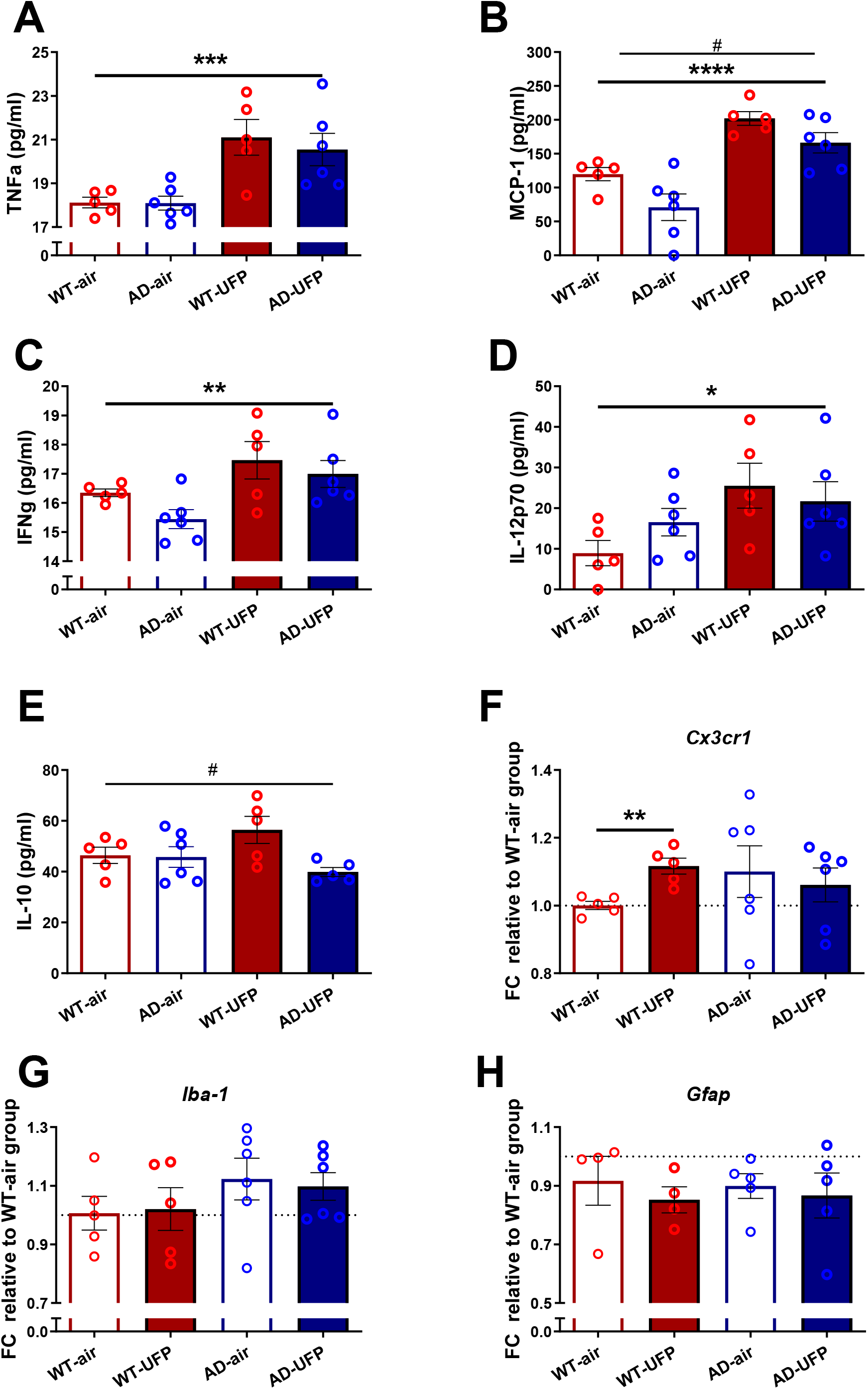
A-E. UFP-induced inflammation in olfactory bulbs of both WT and 5xFAD mice measured by the CBA array after 2-week UFP inhalation exposure. Data are presented as mean (±SEM). Two-way ANOVA was used to measure the genotype and UFP exposure effect between WT and 5xFAD mice. **F-H**. Fold change (FC) of mRNA of glial-specific markers were detected by qPCR in olfactory bulbs of WT and 5xFAD mice after 2-week of inhalation exposure. Data are presented as a mean of a FC relative to WT-air group (±SEM). Nonparametric two-way ANOVA was used to measure the genotype and UFP exposure effect between WT and 5xFAD mice. Because of a significant genotype*UFP interaction, the Mann-Whitney test was used for the comparison of groups within one genotype. «*» UFP inhalation effect: *p < 0.05, **p < 0.01, ***p < 0.001, ****p <0.0001 «#» Genotype effect: #p < 0.05.

To further assess the inflammatory processes induced by UFP exposure in the olfactory bulb, we next analyzed a range of inflammatory markers by qPCR. We observed a slight increase in the microglia-specific fractalkine receptor (*Cx3cr1)* in the olfactory bulbs of UFP-exposed WT (FC=1.12, p < 0.01) but not AD mice (Fig. 2F). However, the mRNA expression of the microglial marker Ionized calcium-binding adaptor protein-1 (*Iba-1*) and glial fibrillary acidic protein (*GFAP*) were unchanged in both treatment groups (Fig. 2G, H).

### 3.4. Inflammatory responses to UFPs in hippocampi

Given that the hippocampus is important for learning and memory and is known to be affected by AD pathology in both humans and 5xFAD mice^47,48^, we next assessed UFP effects in this brain region. Similar to the olfactory bulb, the CBA array revealed that UFP exposure caused alterations in cytokines levels in the hippocampi of both WT and 5xFAD mice as evidenced by increased levels of IFNγ (WT-UFP: 16.93 ± 0.33pg/ml, WT-air: 15.69 ± 0.15pg/ml, AD-UFP:15.42 ± 0.61pg/ml, AD-air: 14.80 ± 0.30pg/ml, p < 0.05) and IL-12p70 (WT-UFP: 19.65 ± 3.34pg/ml, WT-air: 6.612 ± 3.48pg/ml, AD-UFP:11.65 ± 4.13pg/ml, AD-air: 3.58 ± 2.04pg/ml, p < 0.05) after UFP exposure (Fig. 3C, D). Pearson correlation analysis revealed a significant (p < 0.05) positive correlation between the olfactory bulb and hippocampal concentrations of IFNγ (r = 0.49), thus indicating similar responses of this cytokine in both brain areas. Similar to the olfactory bulbs, we observed a higher response to UFP inhalation in the hippocampi of WT mice when compared to AD mice. Genotype-specific differences were found in hippocampal levels of TNFα, MCP-1, IFNγ, and IL-6 (Fig. 3A-C, E).

**Fig. 3.**
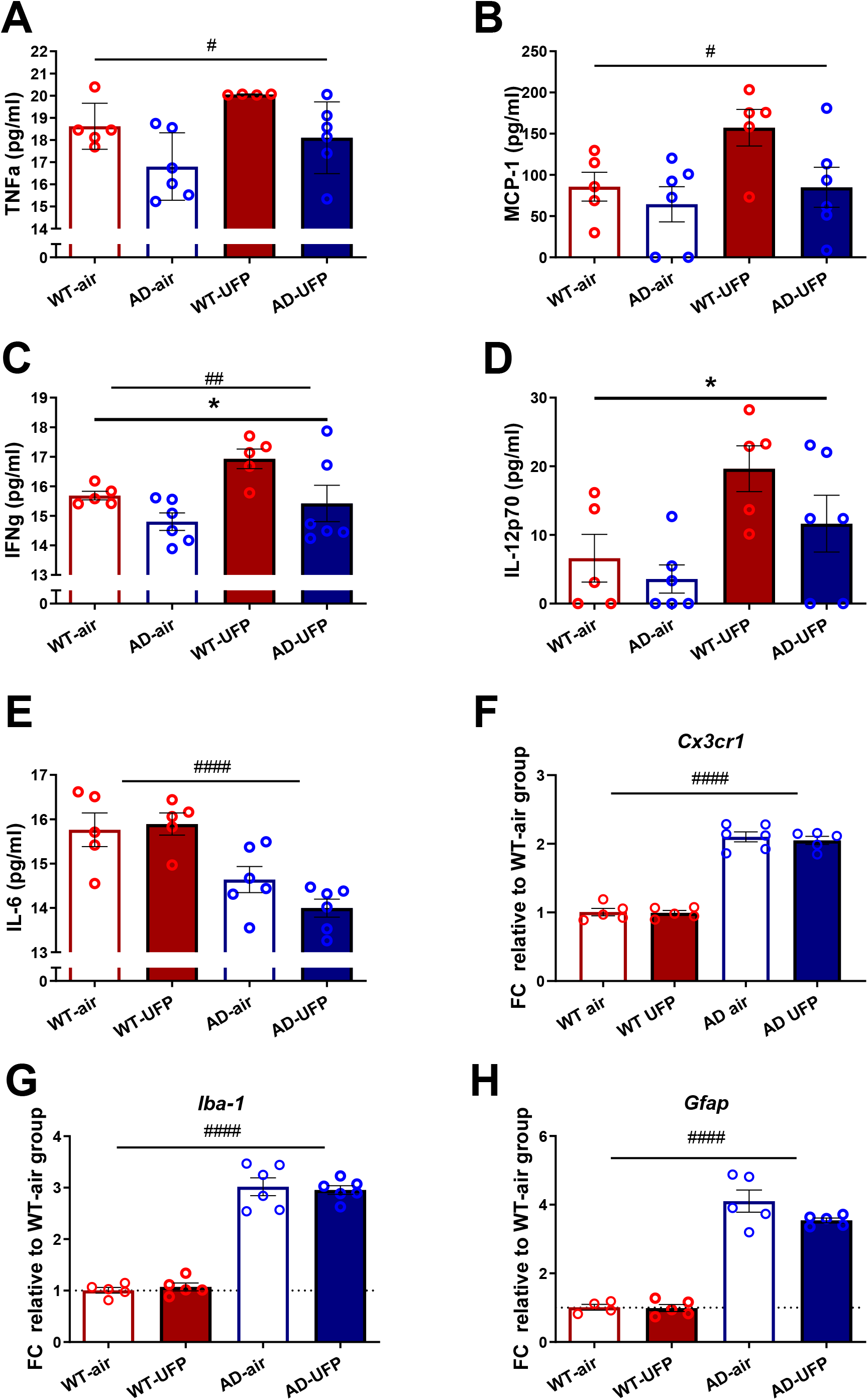
A-E. UFP exposure induced and genotype-specific alterations in inflammatory cytokines in hippocampi of WT and 5xFAD mice after 2-week of inhalation exposure. All data are presented as mean (±SEM). **F-H**. Genotype-specific changes in mRNA levels of Cx3cr1, iba-1, and Gfap in mouse hippocampi. Data are presented as a mean of a fold change (FC) relative to the WT-air group (±SEM). Two-way ANOVA was used to measure genotype and UFP exposure effect between WT and 5xFAD mice. «*» UFP inhalation effect: *p < 0.05, «#» Genotype effect: #p < 0.05 ##p < 0.01, ####p <0.0001.

To complement the CBA data, qPCR was used to assess cell type specific alterations and inflammation-related gene expression changes in the hippocampi of exposed mice. While genotype-dependent increases were observed for the microglial marker *Iba-1* (Fig. 3G) and astrocytic marker *Gfap* (Fig. 3H), the UFP exposure did not influence the expression of these markers. In contrast to the olfactory bulb, hippocampal *Cx3cr1* levels also remained unchanged in the hippocampi after UFP exposure (Fig. 3F). Immunohistochemical stainings for GFAP (Supplementary Fig. 4A) and IBA-1 (Supplementary Fig. 4F) showed an increased population of reactive astrocytes and microglia in the AD hippocampi, as expected. To further examine UFP exposure effects, we quantified the levels of these markers in several brain areas including hippocampus, subiculum, and fimbria fornix (Supplementary Fig. 4). UFP inhalation exposure resulted in a slight, but significant reduction in GFAP protein level specifically in the fornix of 5xFAD mice (AD-UFP: 0.98 ± 0.03%, AD-air: 3.00 ± 0.87%, p<0.05) (Supplementary Fig. 4B). UFP-induced changes were not observed in other brain areas.

### 3.5. Oxidative stress responses to UFPs in hippocampi

Oxidative stress is a widely reported consequence to air pollutant exposure^31,49,50,51,35^. Oxidative stress is commonly reported in pollution exposure studies in both rodent models and epidemiological data as reviewed in ^52^. Previously, chronic exposure studies introduced the hypothesis that the main effect of UFP inhalation is redox imbalance caused by particles that later leads to oxidative stress and neuroinflammation^53,54^ Thus, we analyzed the brains of exposed mice for mRNA expression levels of genes encoding for heme oxygenase-1 (HMOX-1), nuclear factor erythroid 2–related factor 2 (*Nfe2l2*, Nrf2), and NAD(P)H Quinone Dehydrogenase 1 (NQO1). We did not observe a differential gene expression in the olfactory bulbs of UFP-exposed mice, nor was there a genotype-specific difference (data not shown). However, gene expression changes were observed in the mouse hippocampi due to both genotype and inhalation exposure (Fig. 4). The mRNA expression levels of oxidative stress markers were altered in AD mouse hippocampi, as previously reported for *HMOX-1*^55^. In our experiment, mRNA levels of genes of the Nrf2-pathway, including *Nfe2l2* itself and such downstream targets as *Hmox1* and *Nqo1* were significantly up-regulated in the AD hippocampi compared to WT control (Fig. 4A-C). The 2-week inhalation exposure induced a minor up-regulation in the mRNA expression of *Nqo1* in the hippocampi of WT mice (FC=1.07, p < 0.5) (Fig. 4C). After inhalation, a slight reduction was observed for the genes encoding Nrf2 (WT-UFP: FC=0.88 compared to FC=1 for WT-air; AD-UFP: FC=1.52, AD-air: 1.79, p < 0.5) and HMOX-1(WT-UFP: FC=0.89 compared to FC=1 for WT-air; AD-UFP: FC=1.69, AD-air: 1.89, p < 0.01) (Fig. 4A, B). It is important to note that these alterations were less than one-fold, suggesting that the downregulation is relatively minor. Interestingly, Western blot analysis revealed that the level of mitochondrial superoxide dismutase (SOD2) protein was significantly increased upon UFP exposure in the hippocampi of AD by 1.78-fold (p < 0.001), but not in WT mice (Fig. 4D).

**Fig. 4.**
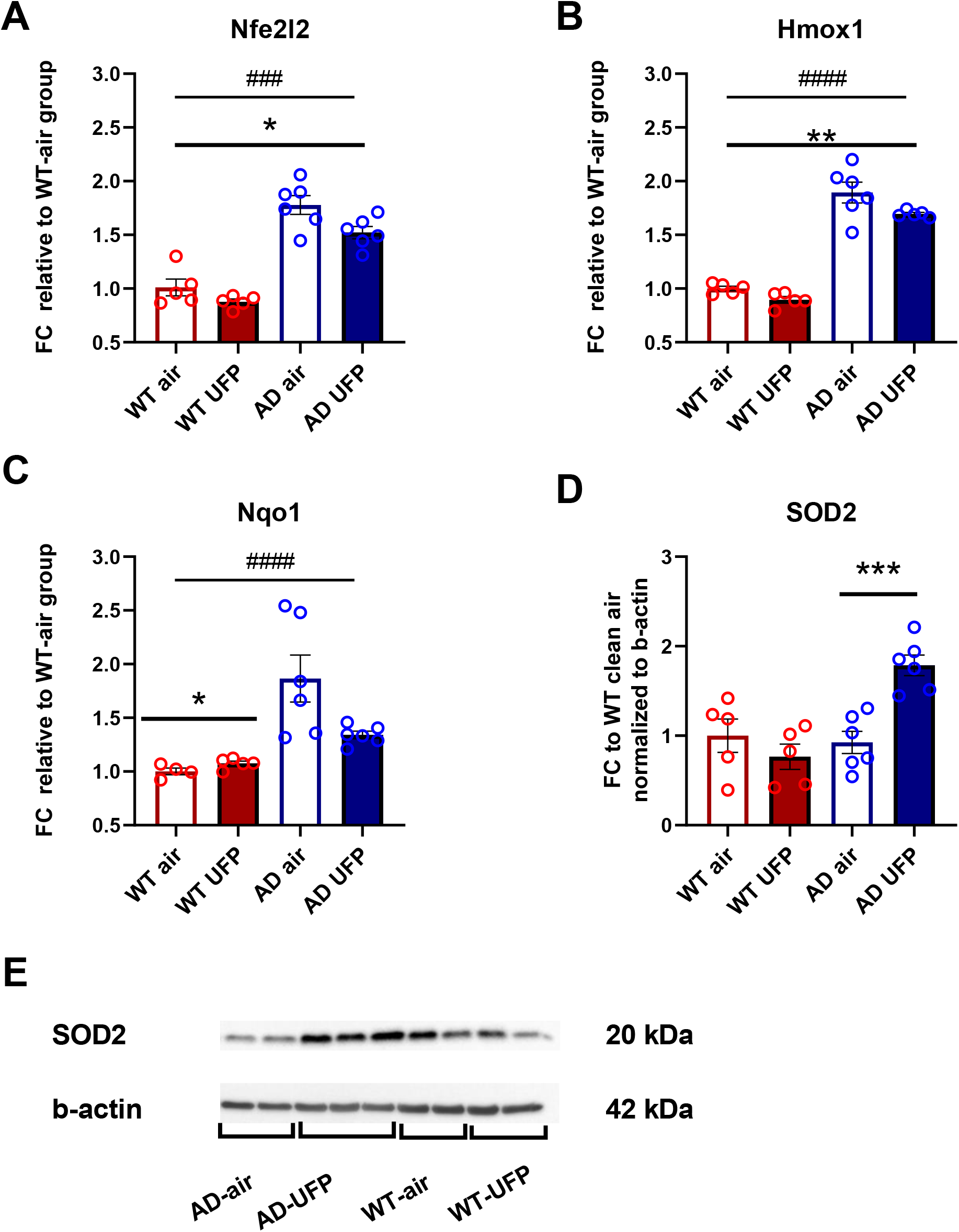
A-C. Hippocampal oxidative stress gene mRNA levels measured by qPCR after 2-week UFP inhalation exposure. All data are presented as a mean of a fold change (FC) relative to the WT-air group (±SEM). Two-way ANOVA was used to measure genotype and UFP exposure effect comparing dCt values in all groups. In case of a significant genotype*UFP interaction, the unpaired t test was used for comparison of groups within one genotype, n = 5-6 per group. «*» UFP inhalation effect: *p < 0.05, *p< 0.01, «#» Genotype effect: ###p < 0.001, ####p < 0.0001. **D**. Western-Blot results for SOD2. Two-way ANOVA was used to measure genotype and UFP exposure effect between WT and 5xFAD mice followed by unpaired t test comparison because of significant interaction between two factors (genotype*UFP). n = 5-6 per group. «*» UFP inhalation effect: ***p < 0.001. **E**. Representative Western-Blot image of SOD2 and b-actin in hippocampal protein samples.

## 4. DISCUSSION

In this study, 6-month-old female 5xFAD and WT mice were exposed to UFPs for 2 weeks through inhalation exposure. Inhalation exposure was chosen as this is the main exposure route to PM in the real-life environment. We focused our study on UFPs due to the ubiquitous nature of this component of air pollution and reported potential for brain translocation^18^. Furthermore, growing evidence supports the harmful effects of the UFPs on brain health in humans and animal models as reviewed in ^13^. It should also be noted that currently there are no legal standards controlling ambient levels of UFPs which indicates an unmet need in changing the ambient air quality standards policy over the globe.

In our exposure design, we used combustion-derived carbonaceous UFPs in a subacute inhalation exposure design with an average of 7.0×10^5^ particles per cm^3^. Assuming that the 5xFAD model lifespan mostly does not exceed more than 2 years of age, a 2-week exposure design represented around 2% of the entire lifespan exposure, which is equivalent to 1.6 years in an 80-year lifespan in humans. The total particle mass concentration inhaled during the study was equivalent to 89.52 µg/m^3^ which corresponds, for example, to the annual average PM2.5 concentrations in countries of South Asia such as India, Bangladesh, and Pakistan, according to air quality index reports between 2019 and 2021 published by IQair agency^56,57^. So far, studies on the toxicological effects of air pollution have mostly focused on diesel exhaust particles (DEP). However, many different combustion sources generate UFPs into ambient air and in general, combustion processes are the major source of ambient air UFPs worldwide. For example, measurements conducted in 2008 in the European Union for anthropogenic outdoor UFPs estimated total UFP emissions to be 271 kilotons in which, for example, road transport contributed to 34% and industrial combustion to 12% of UFP emissions^58^. Therefore, combustion-generated particles are an important source of air pollutants worldwide.

Our study reports findings of UFP exposure in a murine model of AD, which has already been used in other exposure studies with different time-points or exposure design^25,42,59^. In contrast to earlier studies, we did not detect an increase in the brain Aβ load following exposure. That may be explained by the age of animals used in the study which, at 6 months of age, already have extensive amyloid pathology, thus, masking any UFP-induced change in Aβ load. A similar finding was presented in a DEP inhalation study, where increased Aβ plaque load and upregulated brain Aβ42 protein were reported in 4-month-old female 5xFAD mice after 3 weeks of inhalation exposure. However, inhalation-related differences were no longer present at 7 months of age^25^. When accumulation of Aβ was assessed in WT animals, increased levels of Aβ were found in hippocampi of 12-month-old C57BL6/J male mice after inhalation exposure to UFPs^34^. We did not detect any Aβ accumulation in WT littermates of the 5xFAD strain after UFP exposure. This may be explained by both the used exposure design and mouse age.

When the inflammatory response to UFP particles was accessed, in agreement with previous studies, we did not detect UFP-induced changes in most of the measured plasma cytokines. Previously, plasma levels of IL-1b, MCP-1, and G-CSF were reported unchanged in the 5xFAD model after 3 months of exposure to DEP^25^. Our findings reveal that only IFNγ was up-regulated in response to UFP exposure in both genotypes and IL-12p70 in AD-UFP group. From human studies, a link between a polymorphism of IFNγ and fast AD progression was previously demonstrated, indicating an active role of this cytokine in exacerbating the disease^60^. Based on our study results, we postulate that UFP exposure can worsen the preexisting AD pathology by induction of IFNγ. At the time of preparing this manuscript, we did find any studies reporting increased concentrations of IFNγ in rodent plasma after exposure to PM. In humans, however, systemic inflammation is observed after exposure to PM^61^ and increased IFNγ is found in the blood of occupationally exposed persons^62^. These results suggest that this cytokine can serve as a potential biomarker of the systemic inflammatory response in response to UFP exposure in both humans and rodent models and highlights the importance of this cytokine in responses to environmental insults.

The olfactory area is reported to be one of the entry routes of PM to the brain^18^. It was previously reported that UFPs can take the olfactory nerve route, translocate through the olfactory bulb and deposit in both olfactory bulbs as well as in the deeper brain structures such as hippocampus after inhalation^18^,^8^. Previously, an olfaction deficit, accumulation of particles, hyperphosphorylated tau, and inflammation were reported in the olfactory bulbs of Mexico City residents that are constantly exposed to severe air pollution, indicating this organ as one of the main targets of PM effects^24,63^. After two weeks of UFP exposure, we found elevated concentrations of proinflammatory cytokines and chemokines (MCP-1, TNFα, IFNγ, IL-12p70) in olfactory bulbs of both WT and 5xFAD mice. Elevated cytokines levels in the olfactory bulb have been previously reported in mouse inhalation studies with acute inhalation DEP but not UFPs exposure^64^. Our results are in line with these studies, supporting the concept that the olfactory bulbs, which are at the forefront of particle exposure, will show an elevated response when compared to deeper brain areas. To note, we observed a higher magnitude inflammatory response to UFP in WT mice than in AD, indicating the defective immune response of AD mice to the exposure insult. Interestingly, we report that mRNA expression of microglial fractalkine receptor *Cx3cr1*, is elevated in the WT olfactory bulbs after UFP exposure but not in AD mice. Signaling via this receptor is an important regulatory mechanism by which neuronally-derived fractalkine can reduce microglial over-activation during inflammation. The increase in the fractalkine receptor suggests a heightened inflammatory response in the olfactory bulbs of the UFP-exposed mice.

Recently refracted particle deposition was reported to be at the highest level in the hippocampi following chronic inhalation exposure of rats^8^. This suggests that the hippocampus is a target region for PM effects in the brain. Despite the short exposure time used in the current study, we observed hippocampal alterations upon UFP exposure as detected by increased hippocampal IFNγ and IL-12p70 in both WT and AD mice following UFP inhalation. This finding is in line with reports in both chronic and subchronic exposure studies where inflammation has been detected in the hippocampi of WT male mice^7,9,^ and both WT^65, 66^ and AD rats^8^ of both genders after PM exposure^66^. Interestingly, it was reported that inhibition of microglia-derived IL-12 reverses AD-pathology in APP/PS1 transgenic mice, suggesting a crucial role of this cytokine in AD progression^67^. The positive correlation between hippocampal and plasma levels of IL-12 found in the AD group in our study opens up new avenues for research. IFNγ activates microglia and overexpression of this cytokine results in infiltration of peripheral monocytes, suggestive of ongoing inflammation in the hippocampi of the mice used in our study. Interestingly, Pearson’s correlation analysis revealed a significant positive correlation between olfactory bulbs and hippocampi in concentrations of IFNγ in our experiment, therefore indicating the importance of this cytokine in immune responses to UFP in multiple brain areas. The magnitude of the hippocampal inflammatory response was significantly lower than the one observed in the olfactory bulbs, indicating the leading role of this structure located at the forefront of the environmental insult. Summing up the current finding with increased levels of IFNγ in plasma found in our study and increased levels of IFNγ in the blood of occupationally exposed persons^62^, we postulate a key role of this cytokine in systemic and brain responses to air pollution.

While both acute and prolonged exposure studies have previously reported oxidative stress to occur in the olfactory bulbs^64,31^, we did not observe significant UFP-induced alterations in the expression of oxidative stress-related genes. This may, in part be explained by the different genders used in the studies, or the particle composition. While prior acute studies detected robust changes in oxidative stress genes in the olfactory bulbs of male mice, they also used DEP inhalation exposure^64^. We thus believe that both gender and particle composition affect the oxidative stress responses observed in the olfactory bulbs.

When we assessed the effect of UFP exposure in hippocampi, in contrary to previous studies, we found a slight, less than 1-fold reduction in Nrf2-pathway genes that are known to act as an antioxidant defense system of the cell. Interestingly, the nuclear factor κB (NF-κB) protein complex that is known to control cytokine production was previously reported to act as an inhibitor of Nrf2 pathway activation^68^. We hypothesize NF-κB complex activation in our experiment based on increased hippocampal levels of IFNγ after UFP exposure in both genotypes, given that IFNγ is known to act as NF-κB activator^69^. Wakabayashi *et al*., reported that NF-κB can inhibit Nrf2 activity through the recruitment of histone deacetylase to ARE region^70^. Therefore, the observed reduction of the hippocampal Nrf2-pathway genes after UFP-exposure could be due to NF-κB-mediated inhibition. In fact, our correlation analyses revealed a significant inverse correlation between IFNγ concentrations and *Hmox1* mRNA levels in the hippocampi, further supporting our hypothesis (Supplementary figure 5).

When assessing mitochondrial oxidative state, our results demonstrated that the SOD2 protein was significantly up-regulated in the UFP-exposed AD hippocampi. Previously, increased levels of SOD2 have been detected after PM exposure in rat lungs^71^ and in the peripheral blood of non-human primates that were exposed during perinatally development^72^. In the brain, SOD2 mRNA has been shown to be elevated in both hippocampi and striatum of wildtype male rats exposed to UFPs for eight weeks^27^. On one hand, our findings are in agreement with studies linking increased AD pathology with mitochondrial oxidative stress^73^, on the other hand, they refer to a mitochondrial role in PM-induced toxicity as reviewed in ^74, 75, 76^. Taken together, these results indicate that the mitochondrial oxidative state in AD females is more sensitive to UFP effects than that of wildtype littermates.

Our data demonstrate UFP-induced pathological processes in the brains of both AD and WT mice, which supports previous findings (reviewed in^77^), however, some discrepancies were also observed. Importantly, when we compared the concentration of particles with previously published studies that were reviewed in ^78^, it was evident that UFP mass concentrations in our exposure design were relatively low (89.52 compared to up to 350 µg/m^3^ for filter collected UFP and up to 950 µg/m^3^ for DEP nanoparticles reviewed in ^78^). This could very well influence the degree of UFP-induced pathology observed. For example, if we compare the most informative parameter such as the average number concentration of particles per cm^3^ that was around 0.7×10^6^ #/cm^3^ in our study, we found that Hullmann *et al*. reported 2.1×10^6^ in their study where they detected accelerated Aβ plaque formation after inhalation exposure in the same 5xFAD mouse model^25^.For example, we did not detect UFP-induced changes in microglia in the hippocampus even though other papers have shown this ^79,8,80^, and even though altered inflammatory cytokines were observed. It is important also to note that the magnitude of brain responses to PM can depend on the chemical composition of the particles. In our study, we used carbonaceous UFPs, while most often, high magnitude responses have been observed in rodents exposed to DEP. Likewise, the systemic exposure study in mice showed more harmful effects of DEP on the brain in comparison to UFPs, and this was explained by differing chemical composition and reactivity of different types of PM^59^. One feature of our study was that we used a standard soot particle generator, producing a narrow and stable size distribution of ultrafine soot particles. The organic to elemental carbon ratio (OC/EC) of the UFPs generated in this study is on a typical range of real engine exhaust particles^81^. However, real-life engine exhaust particles have always more variation in their composition and particle size distributions, which may partly explain higher magnitude responses in DEP studies. Even though, in most of the cases, DEP studies include very old technology engines and the comparison to real-life is thus impaired. We postulate that more studies comparing different PM particles in one rodent model are needed in order to distinguish the neurotoxicological effects of different types of particles. In our study exposure design, we decided to use a relatively low UFP concentration in order not to exceed the annual average PM concentrations of the countries with the lowest ambient air quality. This allowed us to model the real-life conditions in which humans are regularly exposed to PM concentrations lower than 90 µg/m^3^ throughout their lifespan. However, the fact that with the current exposure design we were able to detect inflammatory responses in plasma, olfactory bulbs, and hippocampi clearly demonstrates that exposure to combustion derived UFPs, even for only two weeks, can have distinct brain health impacts, and together with previous studies findings confirms an urgent need in policy changes for air pollution regulations.

## 5. CONCLUSIONS

In this study, we analyzed and compared the effects of 2-week inhalation exposure to UFPs on the brain and plasma of 6-month-old WT and AD mice. We identified UFP effects on systemic inflammation, neuroinflammation, and oxidative stress induction in the olfactory bulb and hippocampi and addressed genotype-specific implications that were not previously reported in mouse 5xFAD model at this age. Our study differs significantly from previous research with respect to the combination of time of exposure, type of PM, sex of animals and the AD model used for the experiments. Collectively, our findings support epidemiologic studies identifying air pollution as an environmental risk factor and suggest that even relatively short UFP exposure promotes inflammatory mechanisms in the brain. The fact that even a low dose and short-term inhalation exposure already affected the brain’s inflammatory status and oxidative state indicates an urgent need for policy changes controlling UFP levels in ambient air.

## ACKNOWLEDGEMENTS

We would like to thank MSc Valeriia Sitnikova for her help with iba-1 immunostaining and analysis; Mirka Tikkanen for her advises on immunostaining protocols and reagent preparation; MSc Hanna Koponen for the analysis of PM filter samples (gravimetric and thermal-optical).

This study was supported by the Finnish Cultural Foundation, the Academy of Finland, and the Doctoral Program for Molecular Medicine at the University of Eastern Finland.

## ETHICS STATEMENT

The study was carried out in accordance with the Council of Europe Legislation and Regulation for Animal Protection. The Animal Experiment Board in Finland (Regional State Administrative Agency of Southern Finland) approved all the experiments, which were carried out in accordance with EU Directive 2010/63/EU for animal experiments. The study was conducted and reported according to the ARRIVE guidelines (https://arriveguidelines.org/arrive-guidelines/experimental-procedures).

## Notes

### Competing Interest Statement

The authors have declared no competing interest.

## REFERENCES

1. Hahad, O. et al. Ambient Air Pollution Increases the Risk of Cerebrovascular and Neuropsychiatric Disorders through Induction of Inflammation and Oxidative Stress. International journal of molecular sciences vol. 21 4306 (2020).

2. Lee, K. K., Miller, M. R. & Shah, A. S. V. Air pollution and stroke. Journal of Stroke vol. 20 2–11 (2018).

3. Zeng, Y., Lin, R., Liu, L., Liu, Y. & Li, Y. Ambient air pollution exposure and risk of depression: A systematic review and meta-analysis of observational studies. Psychiatry Res. 276, 69–78 (2019).

4. Newbury, J. B. et al. Association of air pollution exposure with psychotic experiences during adolescence. JAMA Psychiatry 76, 614–623 (2019).

5. Qiu, H. et al. Attributable risk of hospital admissions for overall and specific mental disorders due to particulate matter pollution: A time-series study in Chengdu, China. Environ. Res. 170, 230–237 (2019).

6. Peen, J., Schoevers, R. A., Beekman, A. T. & Dekker, J. The current status of urban-rural differences in psychiatric disorders. Acta Psychiatr. Scand. 121, 84–93 (2010).

7. Ehsanifar, M. et al. Learning and memory disorders related to hippocampal inflammation following exposure to air pollution. J. Environ. Heal. Sci. Eng. 1–12 (2021) doi:10.1007/s40201-020-00600-x.

8. Patten, K. T. et al. The Effects of Chronic Exposure to Ambient Traffic-Related Air Pollution on Alzheimer’s Disease Phenotypes in Wildtype and Genetically Predisposed Male and Female Rats. Environ. Health Perspect. 129, EHP8905 (2021).

9. Fonken, L. K. et al. IMMEDIATE COMMUNICATION Air pollution impairs cognition, provokes depressive-like behaviors and alters hippocampal cytokine expression and morphology. Mol. Psychiatry 16, 987–995 (2011).

10. Peeples, L. How air pollution threatens brain health. Proc. Natl. Acad. Sci. U. S. A. 117, 13856–13860 (2020).

11. Haghani, A., Morgan, T. E., Forman, H. J. & Finch, C. E. Air Pollution Neurotoxicity in the Adult Brain: Emerging Concepts from Experimental Findings. Journal of Alzheimer’s disease : JAD vol. 76 773–797 (2020).

12. Kim, H., Kim, W. H., Kim, Y. Y. & Park, H. Y. Air Pollution and Central Nervous System Disease: A Review of the Impact of Fine Particulate Matter on Neurological Disorders. Frontiers in Public Health vol. 8 575330 (2020).

13. Schraufnagel, D. E. The health effects of ultrafine particles. Experimental and Molecular Medicine vol. 52 311–317 (2020).

14. Schmid, O. et al. Dosimetry and toxicology of inhaled ultrafine particles. Biomarkers 14, 67–73 (2009).

15. Kwon, H. S., Ryu, M. H. & Carlsten, C. Ultrafine particles: unique physicochemical properties relevant to health and disease. Experimental and Molecular Medicine vol. 52 318–328 (2020).

16. Xia, T., Zhu, Y., Mu, L., Zhang, Z. F. & Liu, S. Pulmonary diseases induced by ambient ultrafine and engineered nanoparticles in twenty-first century. Natl. Sci. Rev. 3, 416–429 (2016).

17. Nemmar, A. et al. Passage of inhaled particles into the blood circulation in humans. Circulation 105, 411–414 (2002).

18. Oberdörster, G. et al. Translocation of Inhaled Ultrafine Particles to the Brain. Inhal. Toxicol. 16, 437–445 (2004).

19. Livingston, G. et al. Dementia prevention, intervention, and care: 2020 report of the Lancet Commission. The Lancet vol. 396 413–446 (2020).

20. Calderón-Garcidueas, L. et al. Neuroinflammation, hyperphosphorylated tau, diffuse amyloid plaques, and down-regulation of the cellular prion protein in air pollution exposed children and young adults. J. Alzheimer’s Dis. 28, 93–107 (2012).

21. Kilian, J. & Kitazawa, M. The emerging risk of exposure to air pollution on cognitive decline and Alzheimer’s disease – Evidence from epidemiological and animal studies. Biomedical Journal vol. 41 141–162 (2018).

22. Oudin, A. et al. Traffic-Related Air Pollution and Dementia Incidence in Northern Sweden: A Longitudinal Study. Environ. Health Perspect. 124, 306–312 (2016).

23. Crous-Bou, M. et al. Impact of urban environmental exposures on cognitive performance and brain structure of healthy individuals at risk for Alzheimer’s dementia. Environ. Int. 105546 (2020) doi:10.1016/j.envint.2020.105546.

24. Calderón-Garcidueñas, L. et al. Alzheimer’s disease and alpha-synuclein pathology in the olfactory bulbs of infants, children, teens and adults ≤ 40 years in Metropolitan Mexico City. APOE4 carriers at higher risk of suicide accelerate their olfactory bulb pathology. Environ. Res. 166, 348–362 (2018).

25. Hullmann, M. et al. Diesel engine exhaust accelerates plaque formation in a mouse model of Alzheimer’s disease. Part. Fibre Toxicol. 14, 35 (2017).

26. Bhatt, D. P., Puig, K. L., Gorr, M. W., Wold, L. E. & Combs, C. K. A Pilot Study to Assess Effects of Long-Term Inhalation of Airborne Particulate Matter on Early Alzheimer-Like Changes in the Mouse Brain. PLoS One 10, e0127102 (2015).

27. Guerra, R. et al. Exposure to inhaled particulate matter activates early markers of oxidative stress, inflammation and unfolded protein response in rat striatum. Toxicol. Lett. 222, 146–154 (2013).

28. Babadjouni, R. et al. Nanoparticulate matter exposure results in neuroinflammatory changes in the corpus callosum. PLoS One 13, (2018).

29. Aschner, M. & Costa, L. G. Role of inflammation in environmental neurotoxicity.

30. Levesque, S. et al. Diesel exhaust activates and primes microglia: Air pollution, neuroinflammation, and regulation of dopaminergic neurotoxicity. Environ. Health Perspect. 119, 1149–1155 (2011).

31. Calderón-Garcidueñas, L. et al. Long-term air pollution exposure is associated with neuroinflammation, an altered innate immune response, disruption of the blood-brain barrier, ultrafine particulate deposition, and accumulation of amyloid β-42 and α-synuclein in children and young adults. Toxicol. Pathol. 36, 289–310 (2008).

32. Wang, Y., Xiong, L. & Tang, M. Toxicity of inhaled particulate matter on the central nervous system: neuroinflammation, neuropsychological effects and neurodegenerative disease. J. Appl. Toxicol. 37, 644–667 (2017).

33. Costa, L. G. et al. Effects of air pollution on the nervous system and its possible role in neurodevelopmental and neurodegenerative disorders. Pharmacology and Therapeutics vol. 210 107523 (2020).

34. Park, S. J. et al. Exposure of ultrafine particulate matter causes glutathione redox imbalance in the hippocampus: A neurometabolic susceptibility to Alzheimer’s pathology. Sci. Total Environ. 718, 137267 (2020).

35. Milani, C. et al. Systemic exposure to air pollution induces oxidative stress and inflammation in mouse brain, contributing to neurodegeneration onset. Int. J. Mol. Sci. 21, (2020).

36. Stephen, T. L. et al. APOE genotype and sex affect microglial interactions with plaques in Alzheimer’s disease mice. Acta Neuropathol. Commun. 7, (2019).

37. Jawhar, S., Trawicka, A., Jenneckens, C., Bayer, T. A. & Wirths, O. Motor deficits, neuron loss, and reduced anxiety coinciding with axonal degeneration and intraneuronal Aβ aggregation in the 5XFAD mouse model of Alzheimer’s disease. Neurobiol. Aging 33, 196.e29-196.e40 (2012).

38. Sippula, O., Koponen, T. & Jokiniemi, J. Behavior of alkali metal aerosol in a high-temperature porous tube sampling probe. Aerosol Sci. Technol. 46, 1151–1162 (2012).

39. Reagents. ELEMENTAL CARBON (DIESEL PARTICULATE): METHOD 5040, NIOSH Manual of Analytical Methods (NMAM), Fourth Edition. (1999).

40. Ihantola, T. et al. Influence of wood species on toxicity of log-wood stove combustion aerosols: A parallel animal and air-liquid interface cell exposure study on spruce and pine smoke. Part. Fibre Toxicol. 17, 1–26 (2020).

41. Wobbrock, J. O., Findlater, L., Gergle, D. & Higgins, J. J. The Aligned Rank Transform for Nonparametric Factorial Analyses Using Only ANOVA Procedures. (2011).

42. Hullmann, M. et al. Diesel engine exhaust accelerates plaque formation in a mouse model of Alzheimer’s disease. Part. Fibre Toxicol. 14, 35 (2017).

43. Ahn, K. C. et al. Characterization of Impaired Cerebrovascular Structure in APP/PS1 Mouse Brains. Neuroscience 385, 246–254 (2018).

44. Nephew, B. C. et al. Traffic-related particulate matter affects behavior, inflammation, and neural integrity in a developmental rodent model. Environ. Res. 183, 109242 (2020).

45. Licastro, F. et al. Increased plasma levels of interleukin-1, interleukin-6 and α-1-antichymotrypsin in patients with Alzheimer’s disease: Peripheral inflammation or signals from the brain? J. Neuroimmunol. 103, 97–102 (2000).

46. Leung, R. et al. Inflammatory Proteins in Plasma Are Associated with Severity of Alzheimer’s Disease. PLoS One 8, e64971 (2013).

47. Furcila, D., Domínguez-Álvaro, M., DeFelipe, J. & Alonso-Nanclares, L. Subregional Density of Neurons, Neurofibrillary Tangles and Amyloid Plaques in the Hippocampus of Patients With Alzheimer’s Disease. Front. Neuroanat. 13, 99 (2019).

48. Oakley, H. et al. Intraneuronal β-amyloid aggregates, neurodegeneration, and neuron loss in transgenic mice with five familial Alzheimer’s disease mutations: Potential factors in amyloid plaque formation. J. Neurosci. 26, 10129–10140 (2006).

49. Genc, S., Zadeoglulari, Z., Fuss, S. H. & Genc, K. The adverse effects of air pollution on the nervous system. Journal of Toxicology vol. 2012 (2012).

50. Delfino, R. J., Staimer, N. & Vaziri, N. D. Air pollution and circulating biomarkers of oxidative stress. Air Qual. Atmos. Heal. 4, 37–52 (2011).

51. Costa, L. G. et al. Neurotoxicity of traffic-related air pollution. Neurotoxicology 59, 133–139 (2017).

52. Leni, Z., Künzi, L. & Geiser, M. Air pollution causing oxidative stress. Current Opinion in Toxicology vols 20–21 1–8 (2020).

53. Zhang, H. et al. Nrf2-regulated phase II enzymes are induced by chronic ambient nanoparticle exposure in young mice with age-related impairments. Free Radic. Biol. Med. (2012) doi:10.1016/j.freeradbiomed.2012.02.042.

54. Hahad, O. et al. Ambient Air Pollution Increases the Risk of Cerebrovascular and Neuropsychiatric Disorders through Induction of Inflammation and Oxidative Stress. Int. J. Mol. Sci. 21, 4306 (2020).

55. Fernández-Mendívil, C., Arreola, M. A., Hohsfield, L. A., Green, K. N. & Lopez, M. G. Aging and Progression of Beta-Amyloid Pathology in Alzheimer’s Disease Correlates with Microglial Heme-Oxygenase-1 Overexpression. Antioxidants 9, 644 (2020).

56. Empowering the World to Breathe Cleaner Air | IQAir. https://www.iqair.com/world-air-quality-report.

57. 2019 WORLD AIR QUALITY REPORT. https://www.google.com/url?sa=t&rct=j&q=&esrc=s&source=web&cd=&cad=rja&uact=8&ved=2ahUKEwig0s-qjOfwAhXNo4sKHRb9DQAQFjASegQIFBAD&url=https%3A%2F%2Fwww.greenpeace.org%2Fstatic%2Fplanet4-thailand-stateless%2F2020%2F02%2F91ab34b8-2019-world-air-report.pdf&usg=AOvVaw37vxDW-2×79rhb6lltpPsC (2019).

58. European Comission. Industrial emissions of nano- and ultrafine particles - Publications Office of the EU - Final report. Final report https://op.europa.eu/en/publication-detail/-/publication/f5002bc6-ddaa-48cb-9033-a9d12574a32e (2021).

59. Milani, C. et al. Systemic exposure to air pollution induces oxidative stress and inflammation in mouse brain, contributing to neurodegeneration onset. Int. J. Mol. Sci. 21, 3699 (2020).

60. Asselineau, D. et al. Interleukin-10 Production in Response to Amyloid-β Differs between Slow and Fast Decliners in Patients with Alzheimer’s Disease. J. Alzheimer’s Dis. 46, 837–842 (2015).

61. Pope, C. A. et al. Exposure to Fine Particulate Air Pollution Is Associated with Endothelial Injury and Systemic Inflammation. Circ. Res. 119, 1204–1214 (2016).

62. Brucker, N. et al. Biomarkers of occupational exposure to air pollution, inflammation and oxidative damage in taxi drivers. Sci. Total Environ. 463–464, 884–893 (2013).

63. Calderón-Garcidueñas, L. et al. Urban air pollution: Influences on olfactory function and pathology in exposed children and young adults. Exp. Toxicol. Pathol. 62, 91–102 (2010).

64. Cole, T. B. et al. Sex and genetic differences in the effects of acute diesel exhaust exposure on inflammation and oxidative stress in mouse brain. Toxicology 374, 1–9 (2016).

65. Patten, K. T. et al. Effects of early life exposure to traffic-related air pollution on brain development in juvenile Sprague-Dawley rats. Transl. Psychiatry 10, 1–12 (2020).

66. Chang, L. et al. Microglial activation and inflammation caused by traffic-related particulate matter. Chem. Biol. Interact. 311, (2019).

67. Vom Berg, J. et al. Inhibition of IL-12/IL-23 signaling reduces Alzheimer’s diseasea-like pathology and cognitive decline. Nat. Med. 18, 1812–1819 (2012).

68. Liu, G. H., Qu, J. & Shen, X. NF-κB/p65 antagonizes Nrf2-ARE pathway by depriving CBP from Nrf2 and facilitating recruitment of HDAC3 to MafK. Biochim. Biophys. Acta - Mol. Cell Res. 1783, 713–727 (2008).

69. Pfeffer, L. M. The role of nuclear factor κb in the interferon response. Journal of Interferon and Cytokine Research vol. 31 553–559 (2011).

70. Wakabayashi, N., Slocum, S. L., Skoko, J. J., Shin, S. & Kensler, T. W. When NRF2 talks, who’s listening? Antioxidants and Redox Signaling vol. 13 1649–1663 (2010).

71. Stringer, K. A. et al. Particulate phase cigarette smoke increases MnSOD, NQO1, and CINC-1 in rat lungs. Free Radic. Biol. Med. 37, 1527–1533 (2004).

72. Westbrook, D. G., Anderson, P. G., Pinkerton, K. E. & Ballinger, S. W. Perinatal tobacco smoke exposure increases vascular oxidative stress and mitochondrial damage in non-human primates. Cardiovasc. Toxicol. 10, 216–226 (2010).

73. Li, F. et al. Increased plaque burden in brains of APP mutant MnSOD heterozygous knockout mice. J. Neurochem. 89, 1308–1312 (2004).

74. Sharma, J. et al. Emerging role of mitochondria in airborne particulate matter-induced immunotoxicity. Environmental Pollution vol. 270 116242 (2021).

75. Daiber, A. et al. Effects of air pollution particles (ultrafine and fine particulate matter) on mitochondrial function and oxidative stress – Implications for cardiovascular and neurodegenerative diseases. Archives of Biochemistry and Biophysics vol. 696 108662 (2020).

76. Chew, S., Kolosowska, N., Saveleva, L., Malm, T. & Kanninen, K. M. Impairment of mitochondrial function by particulate matter: Implications for the brain. Neurochem. Int. 135, (2020).

77. Costa, L. G. et al. Effects of air pollution on the nervous system and its possible role in neurodevelopmental and neurodegenerative disorders. Pharmacol. Ther. 107523 (2020) doi:10.1016/j.pharmthera.2020.107523.

78. Haghani, A., Morgan, T. E., Forman, H. J. & Finch, C. E. Air Pollution Neurotoxicity in the Adult Brain: Emerging Concepts from Experimental Findings. Journal of Alzheimer’s Disease vol. 76 773–797 (2020).

79. Herr, D. et al. Effects of concentrated ambient ultrafine particulate matter on hallmarks of Alzheimer’s disease in the 3xTgAD mouse model. Neurotoxicology 84, 172–183 (2021).

80. Li, X. et al. Activation of NLRP3 in microglia exacerbates diesel exhaust particles-induced impairment in learning and memory in mice. Environ. Int. 136, 105487 (2020).

81. Pio, C. et al. OC/EC ratio observations in Europe: Re-thinking the approach for apportionment between primary and secondary organic carbon. Atmos. Environ. 45, 6121–6132 (2011).

