## Supplemental Table 1 for "Subacute Inhalation of Ultrafine Particulate Matter Triggers Inflammation Without Altering Amyloid Beta Load in 5xFAD mice"

| *TaqMan primer* | *Catalog number, ThermoFisher Scientific, Waltham, MA, USA* |
| --- | --- |
| *GAPDH* | Mm99999915_g1 |
| *GFAP* | Mm01253033_m1 |
| *Nfe2l2 (Nrf2)* | Mm00477784_m1 |
| *Hmox1 (HO-1)* | Mm00516005_m1 |
| *Nqo1* | Mm01253561_m1 |
| *Aif1 (iba-1)* | Mm00520165_m1 |
| *Gja1 (Cnx43)* | Mm00439105_m1 |
| *Cx3cr1* | Mm00438354_m1 |

**Supplementary Table 1.** *List of TaqMan primers used in this study.*
