## Supplemental Table 2 for "Subacute Inhalation of Ultrafine Particulate Matter Triggers Inflammation Without Altering Amyloid Beta Load in 5xFAD mice"

| *Antibodies for immunohistochemistry* | *Catalog number* | *Dilution* | *Company* |
| --- | --- | --- | --- |
| Amyloid β, clone WO-2, mouse | MABN10 | 1:1000 | Millipore, Burlington, MA, USA |
| Iba-1, rabbit | 019-19741 | 1:250 | Wako, Tokyo, Japan |
| GFAP, rabbit | Z033429-2 | 1:500 | Dako, Glostrup, Denmark |
| GLUT1 antibody, rabbit | ab115730 | 1:500 | Abcam, Cambridge, MA, USA |
| Goat anti-rabbit IgG, Alexa Fluor 488 | A11008 | 1:500 | ThermoFisher Scientific, Waltham, MA, USA |
| Goat anti-mouse IgG, Alexa Fluor 568 | A11004 | 1:500 | ThermoFisher Scientific, Waltham, MA, USA |
| *Antibodies used for Western Blotting* | *Catalog number* | *Dilution* | *Company* |
| β-actin, mouse | A5441 | 1:5000 | Sigma-Aldrich, St. Louis, MO, USA |
| Mn-SOD, rabbit | ADI-SOD-110-D | 1:1000 | Enzo |
| Goat anti-rabbit IgG-HRP conjugate | 170-65-15 | 1:3000 | BioRad, Hercules, CA, USA |
| Donkey anti-mouse IgG-Cy5 cojugate | 715-175-151 | 1:1000 | Jackson Immuno Res. Lab., West Grove, PA, USA |

**Supplementary Table 2.** *List of antibodies used in this study.*
