## Supplemental Figure 1 for "Subacute Inhalation of Ultrafine Particulate Matter Triggers Inflammation Without Altering Amyloid Beta Load in 5xFAD mice"

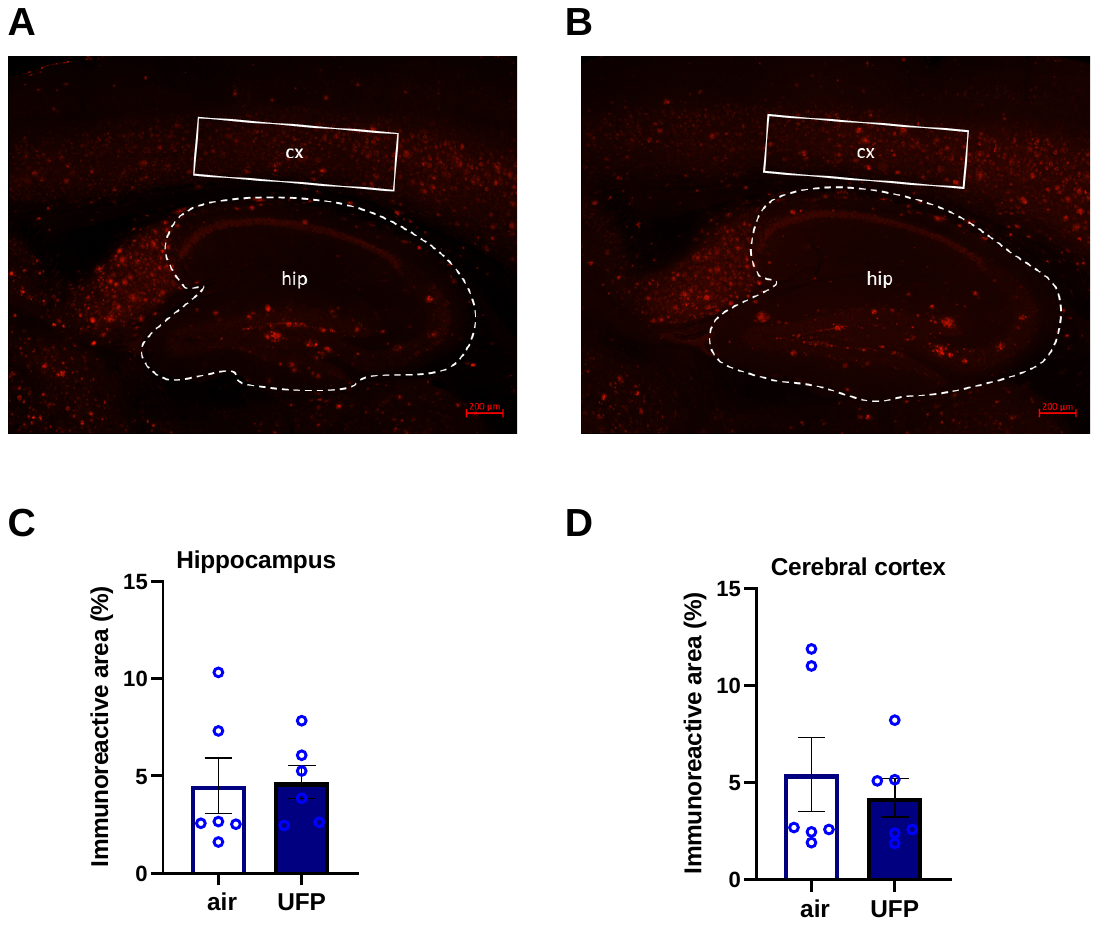

**Supplementary Figure 1**. *Representative images showing* *Aꞵ staining with WO2 antibody in hippocampus (hip) and cortex (cx) of 5XFAD mice (10 x magnification). Representative images of Aβ in AD-air group (****A****, red) and in AD-UFP group (****B****, red).* ***C, D****. Quantification of immunohistochemical staining for Aβ in hippocampi (****C****) and cerebral cortex (****D****) after 2-week inhalation exposure. Protein levels were quantified by measuring the percentage of positive immunoreactive area (%). The Mann–Whitney test was used to measure UFP-exposure effect. All data are presented as mean (± SEM), n = 6 per group.*
