## Supplemental Figure 2 for "Subacute Inhalation of Ultrafine Particulate Matter Triggers Inflammation Without Altering Amyloid Beta Load in 5xFAD mice"

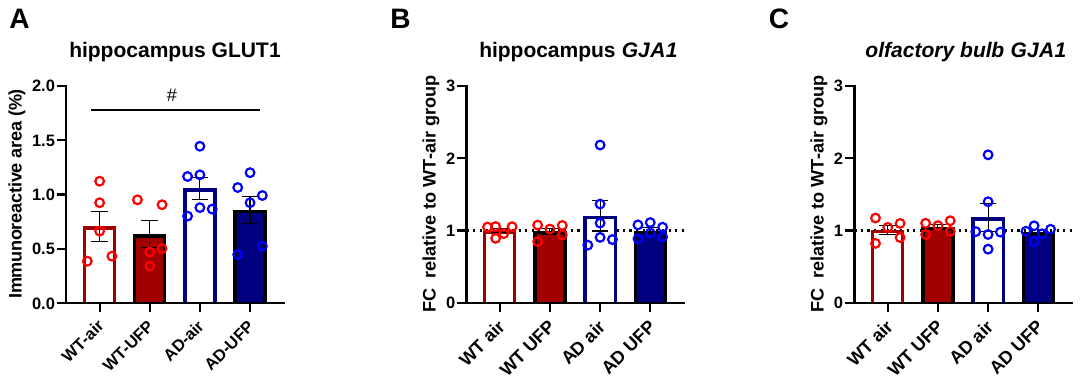

**Supplementary Figure 2. A*.*** *Quantification of immunohistochemical staining for GLUT1 in hippocampi after 2-week inhalation exposure. Protein levels were quantified by measuring the percentage of positive immunoreactive area (%).* *Two-way ANOVA was used to measure genotype and UFP-exposure effect. All data are presented as mean (± SEM). n = 5-6 per group. «#» Genotype effect: # p <0.05.* ***B, C.*** *mRNA levels of GJA1 in hippocampi and olfactory bulbs* *after 2-week inhalation exposure. All data are presented as a mean of a fold change (FC) relative to WT-air group (±SEM). Two-way ANOVA was used to measure genotype and UFP-exposure effect comparing dCt values in all groups.*
