## Supplemental Figure 3 for "Subacute Inhalation of Ultrafine Particulate Matter Triggers Inflammation Without Altering Amyloid Beta Load in 5xFAD mice"

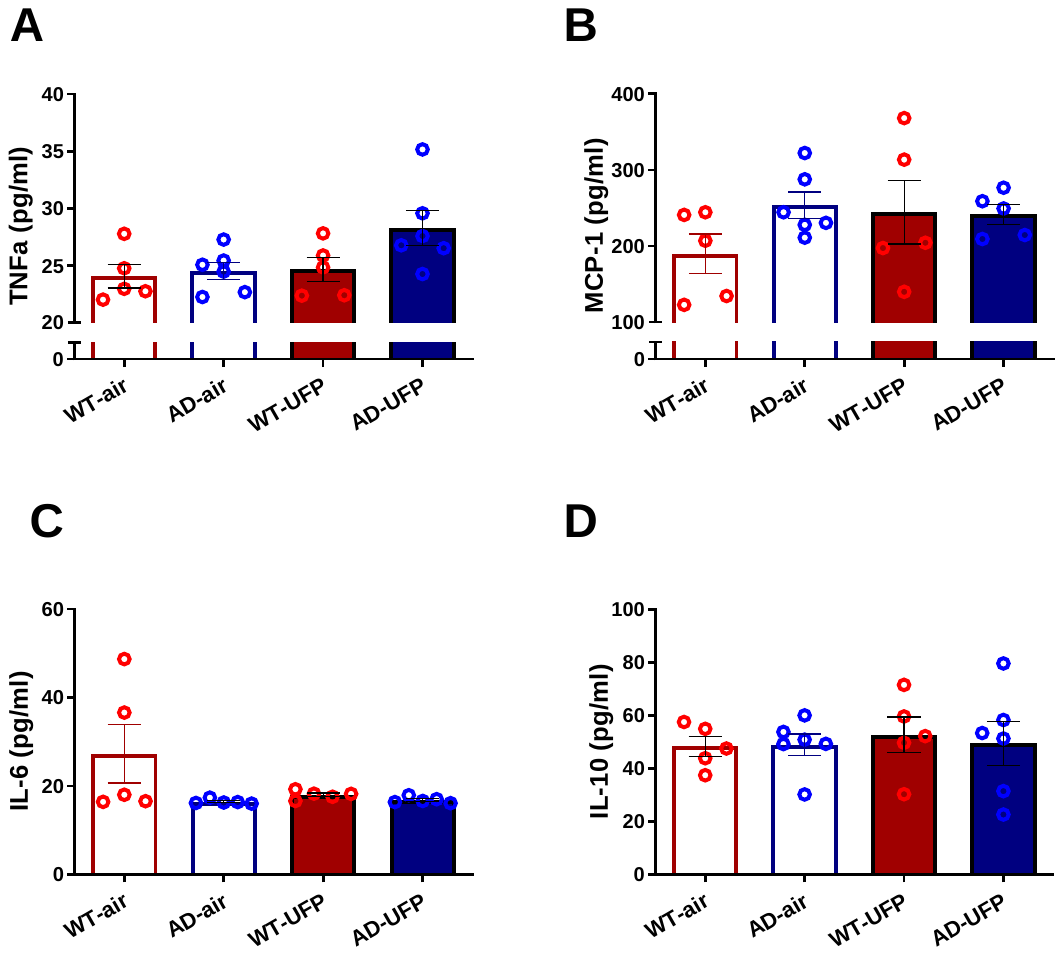

**Supplementary Figure 3***. Plasma cytokines levels after 2-week UFP inhalation exposure. All data are presented as mean (±SEM). Two-way ANOVA was used to measure genotype and UFP exposure effect between WT and 5xFAD mice, n = 5-6 per group.*
