## Supplemental Figure 4 for "Subacute Inhalation of Ultrafine Particulate Matter Triggers Inflammation Without Altering Amyloid Beta Load in 5xFAD mice"

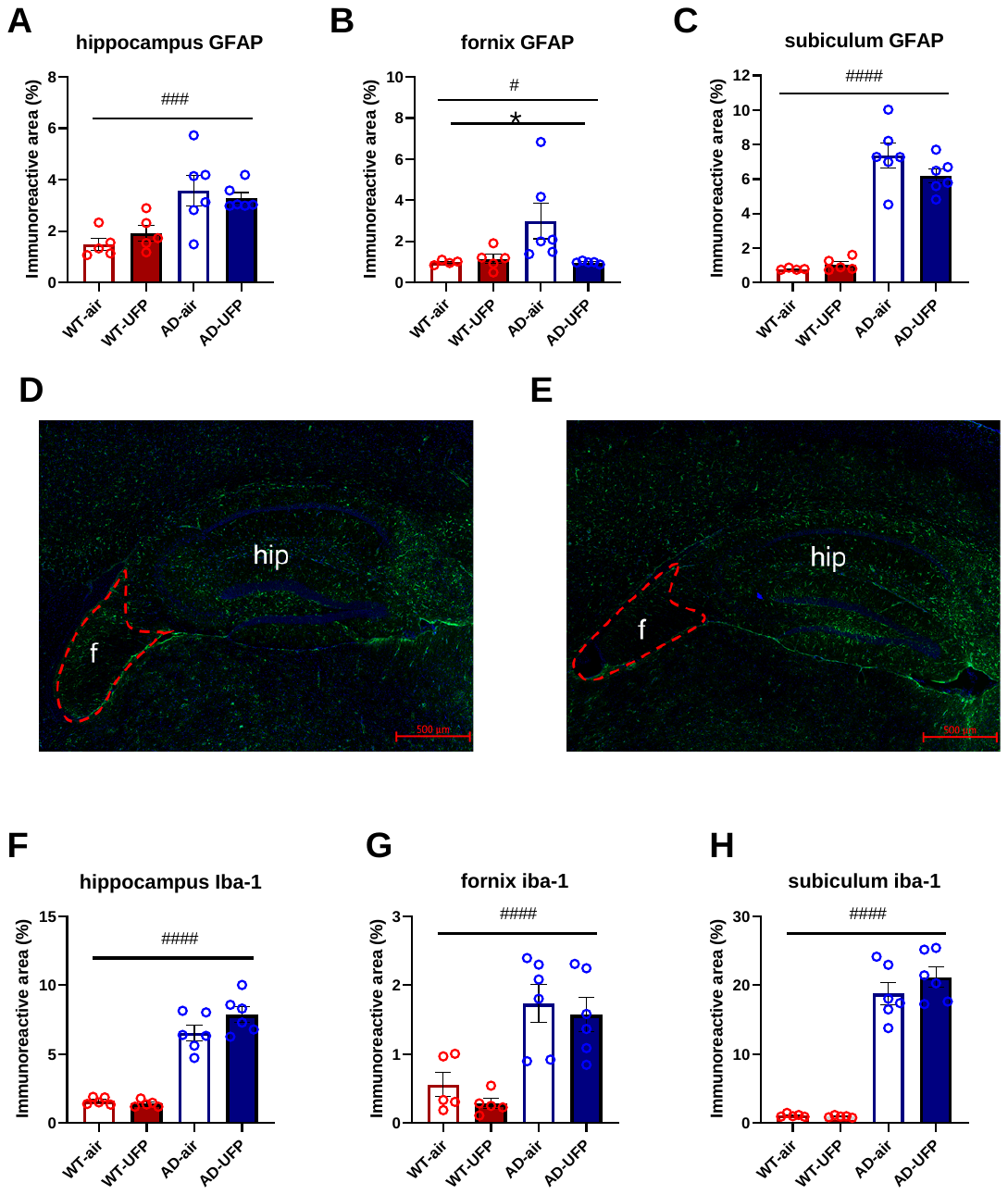
**Supplementary Figure 4.** *Quantification of immunohistochemical staining for GFAP (****4A-C****) and IBA-1 (****4F-H****) in hippocampi, fimbria fornix and subiculum after 2-week inhalation exposure. Protein levels were quantified by measuring the percentage of positive immunoreactive area (%). Two-way ANOVA was used to measure genotype and UFP exposure effect. All data are presented as mean (± SEM). «*» UFP inhalation effect: *p < 0.05, «#» Genotype effect: #p < 0.05, ###p < 0.001, #### p* *<0.0001.* ***D, E****. Representative images showing GFAP (green) and DAPI (blue) staining in hippocampi (hip) and fimbria of fornix (f) of 5XFAD mice (10 x magnification). Representative images of GFAP in AD-air group (C, green) and in AD-UFP group (D, green) after 2-week inhalation exposure.*
