## Supplemental Figure 5 for "Subacute Inhalation of Ultrafine Particulate Matter Triggers Inflammation Without Altering Amyloid Beta Load in 5xFAD mice"

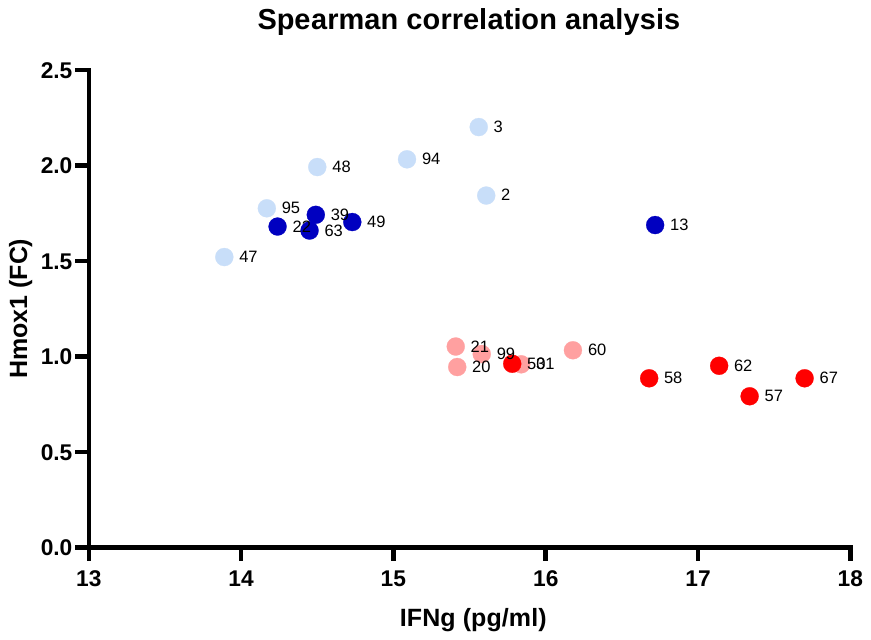

**Supplementary Figure 5.** *Spearman’s correlation between Hmox1 (FC) and IFNg (pg/ml) levels in hippocampi of WT and AD mice after 2-week inhalation exposure showing significant (p < 0.01) inverse correlation (Spearman's correlation coefficient = - 0.60). Light blue dots-AD-air group, dark blue-AD-UFP, light red-WT-air, dark red-WT-UFP; labels – mouse ID numbers.*
